# HiCRep.py: Fast comparison of Hi-C contact matrices in Python

**DOI:** 10.1101/2020.10.27.357756

**Authors:** Dejun Lin, Justin Sanders, William Stafford Noble

## Abstract

Hi-C is the most widely used assay for investigating genome-wide 3D organization of chromatin. When working with Hi-C data, it is often useful to calculate the similarity between contact matrices in order to asses experimental reproducibility or to quantify relationships among Hi-C data from related samples. The HiCRep algorithm has been widely adopted for this task, but the existing R implementation suffers from run time limitations on high resolution Hi-C data or on large single-cell Hi-C datasets. We introduce a Python implementation of HiCRep and demonstrate that it is much faster than the existing R implementation. Furthermore, we give examples of HiCRep’s ability to accurately distinguish replicates from non-replicates and to reveal cell type structure among collections of Hi-C data. HiCRep.py and its documentation are available with a GPL license at https://github.com/Noble-Lab/hicrep. The software may be installed automatically using the pip package installer.

## 1 Introduction

Hi-C is a powerful genomic assay for quantifying chromatin interactions across the whole genome [1]. It has been used extensively to study genome architecture and function in many different species and to understand how genome structure affects genetic diseases. The result of a Hi-C experiment is typically processed into a matrix, whose entries are contact counts between pairs of genomic loci. As Hi-C experiments become more popular, tools that are able to efficiently perform analysis on the resulting contact matrices are in increasing demand [2].

A common task in Hi-C data analysis is measuring the similarity between pairs of data sets. One application of Hi-C similarity is to assess experimental reproducibility. Low reproducibility may indicate low experiment quality or low sequencing depth. Low reproducibility may also warn against merging multiple Hi-C replicates, which is a common practice to boost the signal-to-noise ratio [3]. HiCRep is a tool for quantifying the similarity between pairs of Hi-C contact matrices based on their stratum adjusted correlation coefficients (SCCs) [4]. The SCC is a correlation score ranging from − 1 and 1, where a higher score suggests higher similarity between the two input Hi-C matrices. Using high SCC scores as a proxy for high reproducibility, a number of published works have used HiCRep to asses the quality of replicate experiments and to make sure that merging them is sound [5, 6] or to validate that data from a novel assay closely resembles traditional Hi-C data [7, 8]. Beyond comparing replicates, HiCRep has also proved useful as a tool for measuring quantitative differences among samples. For example, Ray et al. used HiCRep to compare Hi-C contact maps of samples before and after undergoing heat shock in order to determine whether the shock had an effect on chromatin structure [9]. HiCRep can also be used to help interpret single-cell Hi-C (scHi-C) data. For example, Liu et al. demonstrated that the SCC values calculated by HiCRep can be used as the basis for a multidimensional scaling (MDS) visualization that accurately captures cell cycle structure in scHi-C data [10].

The original implementation of HiCRep was released as an R package [4]. One of its the biggest drawbacks is its inefficiency, mainly because of the dependence on dense contact matrix operations. In a head-to-head comparison against three other tools for measuring reproducibility, HiCRep was found to be the slowest by a significant margin [3]. This means that applying the R implementation to Hi-C data at high resolution or to large scHi-C data sets is prohibitively slow.

Here we present a Python implementation of the HiCRep algorithm that is much faster than its predecessor. Our Python version implements all operations using sparse matrices, which greatly reduce the memory consumption and computation time. Additionally, we have made efforts to make the software more accessible by providing a command line interface as well as a Python application programming interface.

## 2 Implementation

HiCRep takes as input two Hi-C contact matrices in either.cool or.mcool format [11]. First, matrices are normalized by the total contact counts and smoothed with a 2D mean filter of size set by the user. Then, corresponding diagonals of the two contact matrices are compared and used to calculate SCC scores, as described in the original HiCRep paper [4]. The software produces as output a list of of SCC scores per chro-mosome. This output faithfully matches that produced by the existing R implementation (see Supplement for details). We provide thorough unit tests of the implementation covering most of its functionality.

## 3 Results

We used HiCRep to calculate SCC scores between 95 pairs of publicly available Hi-C matrices—19 pairs of biological replicates, 38 pairs of non-replicates of the same cell type, and 38 pairs of non-replicates of different cell types. As shown in Figure 1A, pairs of replicates consistently exhibit very high SCC scores (mean: 0.98, stdv: 0.02), which are markedly higher than the scores of both non-replicates of the same cell type (mean: 0.86, stdv: 0.10) and non-replicates of different cell types (mean: 0.61, stdv: 0.16). These results suggest that HiCRep does a good job of capturing the reproducibility of Hi-C datasets and is able to accurately separate replicates from non-replicates.

**Figure 1:**
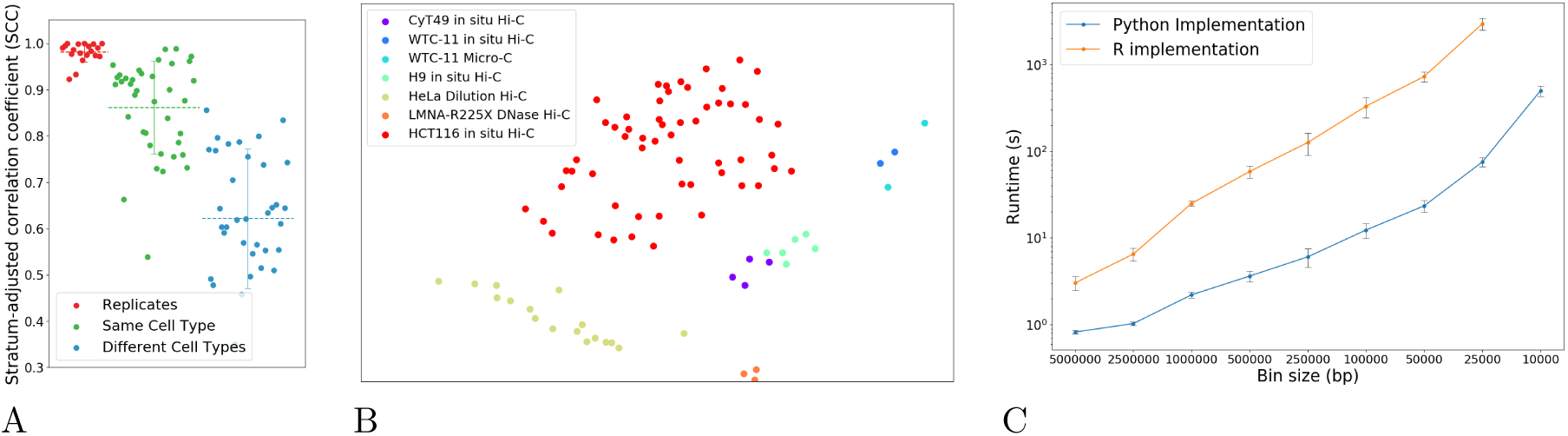
**(A)** The figure plots the HiCRep score for various pairs of Hi-C experiments, including biological replicates (red), non-replicate experiments of the same cell type (green), and non-replicate experiments of different cell types (blue). Horizontal lines with error bars correspond to the mean and stdev of each group. **(B)** Multidimensional scaling plot based on HiCRep scores from 90 Hi-C experiments carried out on a variety of cell types. **(C)** Timing comparison of the R and Python implementations of HiCRep for Hi-C matrices with varying bin sizes. Error bars are standard deviation over five runs.

We also used HiCRep to evaluate the pairwise SCC scores of 90 Hi-C experiments conducted by the 4D Nucleome Consortium on a number of different cell types (Supplementary Table 1). Using these SCC scores as the distance metric for an MDS model, we show that HiCRep reveals structure among the experiments, with different cell types clustering separately (Figure 1B).

Finally, we compared the run times of our implementation of HiCRep to the R implementation. We selected five pairs of high resolution Hi-C experiments and ran both implementations of HiCRep on each of them at a number of different resolutions (Figure 1C). Comparing the runtimes, we see that at higher resolutions the Python implementation of HiCRep is more than 20 times faster than the R version. This speed increase allows our version of HiCRep to be practically applied to data with much smaller bin sizes or to larger collections of scHi-C data than was previously possible.

## Supplement

### Verifying the reimplementation

In order to to ensure that our Python implementation reproduces the results of the R version, we ran both versions on five pairs of Hi-C data sets at different resolutions and with different parameters, and verified that the resulting SCC scores matched perfectly. We also include in our implementation a comprehensive set of unit tests that ensure critical functions are working as expected.

### Data

All of the Hi-C contact matrices presented here were obtained from the 4DN data portal, where they are available in Cooler format (https://data.4dnucleome.org). A list of accession numbers for the files we used are in Supplementary Table 1. No normalization was performed on any of the contact matrices.

### Comparison of replicates

To assess how well HiCRep can separate replicates form non-replicates, we downloaded 19 pairs of replicates from 4DN. To produce Figure 1A, we used HiCRep to compute SCC scores for the 19 replicate pairs as well as 38 randomly selected pairs of data sets from the same “biosource” (i.e., cell type or tissue sample) and 38 randomly selected pairs from different biosources. HiCRep was run with a bin size of 500 kb, a smoothing factor *h* = 2, a maximum genomic distance of 5 Mb, and down-sampling set to false.

### Multidimensional scaling

The MDS plot in Figure 1B was produced by using HiCRep to obtain pairwise SCC scores for 90 Hi-C datasets where replicates had already been merged. We then converted these correlation coefficients to Euclidean distance with the equation 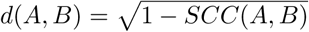. Finally, these distances were passed as input to the scikit-learn implementation of MDS [12] called with maximum iterations set to 30,000 and number of initializations set to 10. For this analysis, HiCRep was run with the same settings as listed above.

### Timing

For the timing experiment, both versions of HiCRep were run on a computer with an Intel i7 CPU at 1.10 GHz with 16GB of RAM. Our Python implementation of HiCRep was installed from PyPI and run with Python 3.7.6, NumPy 1.18.5, and SciPy 1.5.1, all of which were installed through the conda package manager. The R implementation was installed from conda’s bioconda channel and run with R version 4.0.2, also installed from conda. In order to ensure a fair comparison, we undertook measures to ensure that NumPy was not parallelizing any operations and that R was not forced to use swap space.

